# Bound-state diffusion due to binding to flexible polymers in a selective biofilter

**DOI:** 10.1101/736942

**Authors:** L. Maguire, M. D. Betterton, L. E. Hough

## Abstract

Selective biofilters are used by cells to control the transport of proteins, nucleic acids, and other macromolecules. Biological filters demonstrate both high specificity and rapid motion or high flux of proteins. In contrast, high flux comes at the expense of selectivity in many synthetic filters. Binding can lead to selective transport in systems in which the bound particle can diffuse, but the mechanisms that lead to bound diffusion remain unclear. Previous theory has proposed a molecular mechanism of bound-state mobility based only on transient binding to flexible polymers. However, this mechanism has not been directly tested in experiments. We demonstrate that bound mobility via tethered diffusion can be engineered into a synthetic gel using protein fragments derived from the nuclear pore complex. The resulting bound-state diffusion is quantitatively consistent with theory. Our results suggest that synthetic biological filters can be designed to to take advantage of tethered diffusion to give rapid, selective transport.

**SIGNIFICANCE:** Biological filters control the passage of proteins and other macromolecules between compartments of living systems. Determination of molecular mechanisms giving selective transport would enable the design of both selective filters and particles designed to penetrate biological barriers for drug delivery. One such mechanism arises from transient binding to dynamic polymer tethers. We designed a biomaterial which supports this type of tethered diffusion, demonstrating the potential to engineer bio-inspired filters.

## INTRODUCTION

Living systems depend on molecular filters in order to selectively control transport of macromolecules. Binding interactions affect selective filtering in a wide range of biological systems, but the molecular mechanisms by which binding leads to selectivity are often unclear. In some cases, particles which bind to a selective barrier are hindered while inert particles pass more readily. For example, nanoparticles designed for drug delivery through mucosal membranes are most effective when their binding interactions to mucus are minimized (1–3). In other cases, biofilters use binding to enhance the flux of transported proteins, as in the case of the nuclear pore complex (NPC). Proteins which bind to the barrier have a higher flux through it than do inert proteins (4–6). This suggests that the function of binding in selective biological filters is complex and context-dependent, motivating further study.

The nuclear pore complex is a filter which relies on binding to control protein flux. The selective barrier of the NPC controls transport between the cytoplasm and nucleus. It consists of a channel approximately 50 nm in diameter and 100 nm long which is filled with intrinsically disordered FG nucleoporins (FG Nups), named for their many short hydrophobic phenylalanine-glycine (FG) motifs in an otherwise hydrophilic protein (Fig. 1A) (7, 8). Transport factors are proteins that bind to the FG motifs and pass rapidly through the NPC, carrying cargo with them. In contrast, non-binding proteins larger than about 30 kDa have much lower flux through the pore than similarly-sized transport factor-cargo complexes (4–6). While models of diffusion within the NPC have been developed to explain selectivity, many reduce the problem to that of a particle diffusing within an effective energy landscape (6, 9–12), an approach which does not address the molecular mechanisms of diffusion that could lead to selectivity. Therefore, we sought to investigate mechanisms of binding and mobility that can lead to selective transport.

**Figure 1:**
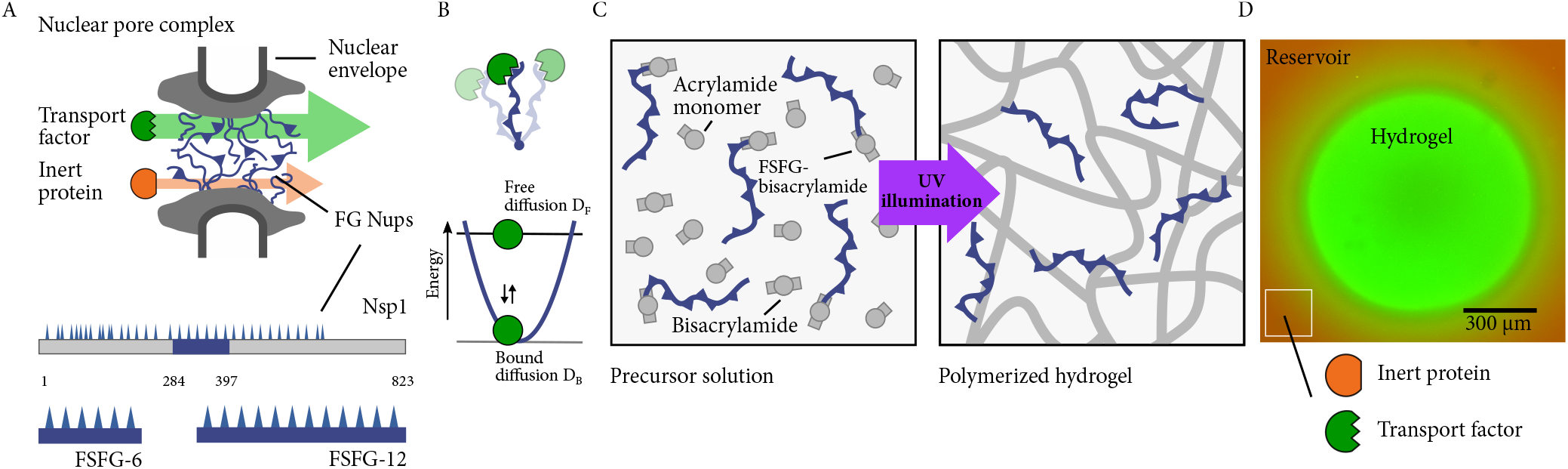
(A) Schematic of nuclear pore complex and FG constructs. The nuclear pore complex (grey) is filled with FG Nups (blue polymers) that selectively passage transport factors (green) that bind to FG Nups while blocking non-binding proteins (red). FSFG_6_ and FSFG_12_ are peptides derived from the FG Nup Nsp1 containing 6 or 12 FSFG motifs. (B) Schematic showing that transport factors bound to an FG Nup fragment can retain mobility (top), and the energy landscape used to model a protein undergoing free diffusion (flat line) or bound diffusion (harmonic well) when bound to a flexible tether (bottom). (C) Schematic of gel fabrication. A precursor solution is polymerized upon exposure to UV illumination, tethering FG peptides to the resulting gel. (D) An experimental image of a gel equilibrated in a reservoir containing a fluorescent transport factor and inert protein.

Recent work has highlighted the importance of bound mobility in selective transport (13–15). A particle’s steady-state flux through a filter can be increased by binding if the bound particle remains mobile (16–18). An open question is what molecular mechanisms lead to this mobility. Previous work made the intuitive proposal that a filter can sustain bound mobility if all of the components, including the binding motifs, are mobile (16). However, in a filter, the attachment site for a binding motif typically is stationary. In this case, several potential mechanisms can contribute to motion of a particle while bound. For example, if binding motifs are tightly spaced, then particles with multiple binding sites can move between binding motifs without unbinding (14, 15). Another possible molecular mechanism for bound diffusion is diffusion during short-lived binding interactions with flexible polymers (13). Dynamic polymers allow a bound particle to move and may additionally drive motion due to elastic kicks when extended tethers bind (Fig. 1B) (13, 19, 20). When applied to transport factors binding to disordered FG Nups within the NPC, a theory based on tethered diffusion agrees with experimental measurements of selective transport of NTF2 (4, 13, 21, 22) suggesting that this mechanism may contribute to selective transport through the nuclear pore complex.

## MATERIALS AND METHODS

### Protein expression, purification, and labeling

We aimed to to use proteins with rapid dynamics, including diffusion-limited on-rates, and well-established binding affinities. For this reason, we chose yeast nuclear transport factor 2 (NTF2) and synthetic fragments derived from the yeast FG Nup Nsp1 (FSFG_6_ and FSFG_12_ (23, 24)). NTF2 is of similar size to the red fluorescent protein mCherry, which is inert to the FG-Nup-based protein fragments. NTF2, FSFG_6_, FSFG_12_ and mCherry (a gift from Amy Palmer) were expressed in BL21 DE3 Gold cells. All proteins contained a C-terminal his tag. FG constructs also contained a terminal cysteine. Cultures were grown in LB to OD 0.6-0.8 and then induced for 2-4 hours with 1 mM IPTG (NTF2 and FG constructs) or 100 mM IPTG overnight (mCherry). Periplasmic matrix was removed by resuspending cell pellets in SHE buffer (20% sucrose, 50 mM HEPES, 1 mM EDTA), spinning down and resuspending in 5 mM MgSO_4_, incubating 10 minutes on ice, and spinning down and discarding supernatant (25). Proteins were purified with a cobalt affinity column in potassium transport buffer (PTB; 150 mM KCl, 20 mM HEPES, 2 mM MgCl_2_).

To conjugate the FG constructs to the gel scaffold, we attached a reactive bisacrylamide moity to the thiol of the FG constructs. The protein was reduced using immobilized TCEP reducing gel (Thermo Fisher) in a spin column and spun into a solution of 300 mM triethanolamine and 10-fold molar excess bisacrylamide. Reaction mixture was incubated shaking at room temperature for 30 min, dialyzed into 25 mM NH_4_HCO_3_, and lyophilized. The labeling efficiency was quantified with an Ellman’s reagent assay.

For visualization, NTF2 was labeled with fluorescein-NHS. NTF2 in PTB was added to 15-fold molar excess fluorescein-NHS and incubated stirring at room temperature for 1 hour. Reaction mixture was re-purified using a cobalt affinity column, washing with at least 100 column volumes of PTB prior to elution to remove unreacted dye. Labeled NTF2 was eluted with 300 mM imidazole in PTB, dialyzed into PTB to remove imidazole, and frozen within 24 hours of labeling to minimize dye hydrolysis. Labeled NTF2 was thawed immediately before use.

### Gel fabrication

Polyacrylamide gels were chosen as the substrate due to the ease of tuning their properties and their biocompatibility (26–29). Gel precursor solutions were prepared with 6% v/v acrylamide and 0.2% v/v bisacrylamide (30% acrylamide/bisacrylamide stock 29:1, BioRad), 2 mM LAP photoinitiator (Sigma) (30), and 0.67 mM resuspended FSFG-bis in PTB (10 mg/mL FSFG_6_ or 20 mg/mL FSFG_12_). Precursor solutions were protected from light and degassed for 10 minutes immediately before polymerization.

We designed a flow chamber using a PDMS mold to allow for facile gel fabrication and exchange of the surrounding solution. The chambers were approximately 400 *μ*m deep, chosen so that the majority of a 0.5 or 1.0 *μ*L polymerized droplet fit within the field of view of a 4x objective. Unpolymerized precursor solution was placed on a coverslip. The chamber was closed, and the gel was polymerized by a 30-second exposure to 365-nm light at approximately 220 mW/cm^2^. In order to remove any unconjugated precursor, gels were then rinsed with 100 gel volumes of PTB and soaked in PTB overnight at 4°C inside the sealed flow chambers.

### FRAP measurements

In order to measure the bound diffusion of NTF2 and mCherry, we measured the diffusion constant within the gel (an average over both bound and free motion). FRAP is a well established method for measuring diffusion, and provided sufficient precision for our needs (31). Fluorescence recovery after photobleaching (FRAP) was performed on gels that had been equilibrated with NTF2 and mCherry. Wash buffer was removed by pipette from the flow chamber reservoir and replaced with 20 *μ*M NTF2-fluorescein and 20 *μ*M mCherry in PTB. The chamber was resealed and equilibrated for 24 hours. Photobleaching was then performed with an Olympus IX-81 widefield microscope with a Prior Lumen 200 Metal-Halide lamp. First, a reference image was taken at 4x magnification. A circular region of the gel approximately 300 *μ*m in radius was then photobleached using a 5-second exposure at full power through a DAPI (352-402 nm excitation / 417-477 nm emission) filter cube at 40x magnification. Following the bleach, the 4x objective was rapidly returned and a time series recorded in FITC and TRITC channels with a Hamamatsu ORCA-ER C4742-80 camera. A typical series consisted of 15-30 frames recorded as rapidly as possible (5-10 s per frame), followed by 30-60 frames recorded at 1-2 minutes per frame. The total experiment time was 1-4 hours. Typical exposure times were 10 ms (FITC) and 40 ms (TRITC), both with a gain of 3 dB.

The fluorescence recovery curve was created by calculating the average intensity in the bleach spot as a function of time and dividing that value by the average intensity of the entire gel over time. This normalization method compensates for photobleaching during recovery, as verified by simulating data with varying photobleaching rate.

### Fluorescence recovery curve fitting

FRAP analysis of systems with both diffusion and binding differs based on the relative timescales of diffusion and reaction kinetics. Three basic regimes have been identified: diffusion-dominant, effective-diffusion, and reaction-dominant (32, 33). Our system falls into the effective-diffusion regime in which the reaction is fast relative to diffusion. In this case, a pure diffusion model can be used with an effective diffusion constant. Previously-developed models do not account for continual exchange between the gel and surrounding reservoir (33–36). As this exchange is significant over the timescale of our experiments, we modeled fluorescence recovery using a full time-dependent Fourier series solution to the diffusion equation.

We solved the diffusion equation 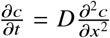 in a circular region of radius *a*, centered at the origin, whose boundary is held fixed at concentration *c* = 0. The first post-bleach image is used as the initial concentration distribution *f* (*r*, *θ*) in the region *r* < *a*. Green’s function integrals for the appropriate boundary conditions (37) allow the concentration *c*(*r*, *θ*, *t*) to be calculated for *r* < *a*. The full solution to the diffusion equation within the gel is given by

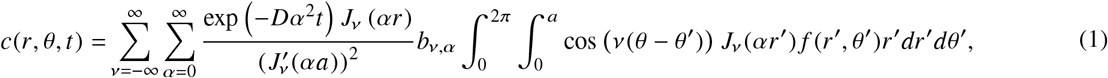

where (*αa*) are the zeros of the Bessel function of the first kind *J*_*ν*_, and *b*_*ν*,*α*_ are weighting constants described below. The sums run over all positive and negative integer Bessel orders and over all zeros of each Bessel function. The boundary is held at *c* = 0, so it must be shifted by an offset in order to match the experimental concentration at the boundary, taken to be just within the gel. The area of the gel visible within the microscope image is denoted Ω. Rewriting Eq. 1 to remove the unprimed coordinates from the integral, the mode coefficients *C*_*ν*,*α*_ and *S*_*ν*,*α*_ can then be defined as

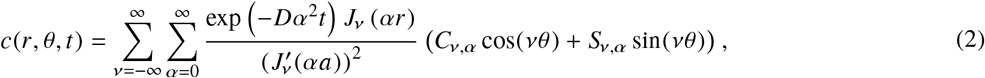

with

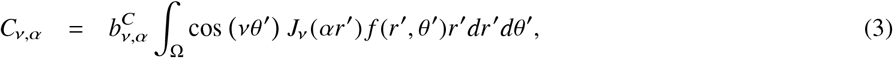

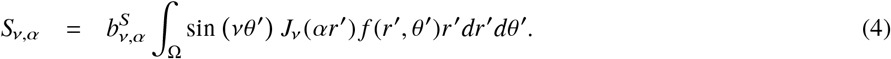

The weighting constants *b*_*ν*,*α*_ would be equal to 2/*πa*^2^ for every mode if the entire gel were within the field of view of the microscope, which is not always the case. To compensate for the area outside of the field of view, they are instead given by

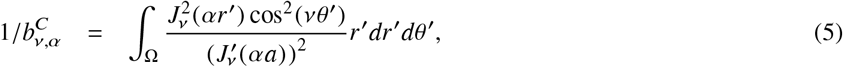

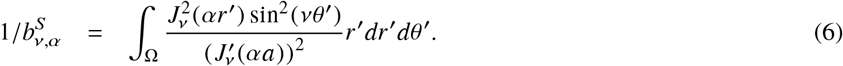

The mode coefficients were calculated numerically. The polar coordinates (*r*, *θ*) were converted to Cartesian (*x*, *y*) and a sum taken over all the pixels of the initial post-bleach image *f* (*x*, *y*) within Ω. The origin was set at the center of the gel and *a* was estimated numerically. Before calculating the mode coefficient, the average intensity of the equilibrated portions of the gel was subtracted from the entire image, effectively setting the zeroth order coefficient to zero. The sum was scaled using the area per pixel and normalized using the weighting constants *b*_*ν*,*α*_.

Once the mode coefficients were calculated, the series solution was constructed using Eq. 2. The average intensity of the bleach spot over time 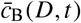 was determined by integrating the series solution over the bleach spot:

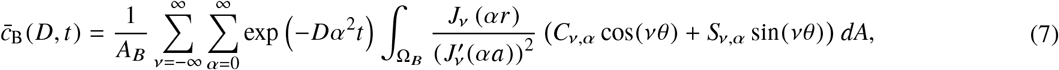

where Ω_*B*_ is the bleach spot and *A_B_* is its total area. The average intensity of the gel 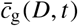 was determined similarly. The equilibrium concentration *c*_0_ was added back to both and the two values divided to represent the normalized intensity of the bleach spot over time. Two additional parameters, *c*_1_ and *c*_2_, were incorporated to reflect the bleach depth and final recovered concentration. Finally, the recovery curve was fit to the equation

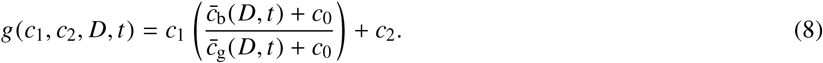

The values of *c*_1_ and *c*_2_ were of order unity and were not used in further analysis. The diffusion constant *D* was used to determine the bound diffusion constant as described below.

### Partition coefficient analysis

To calculate a protein’s bound diffusion constant within a gel, we determined both the diffusion constant within the gel and the fraction of time the protein is bound. The partition coefficients of NTF2 and mCherry (Fig. 2) can be used to calculate the fraction of time that NTF2 spends bound within the FG gels. When the system is in equilibrium, the concentration of free transport factor (*T*), free Nup (*N*), and transport factor - Nup complex (*C*) is related to the dissociation constant *K_D_* by *K_D_* = *NT*/*C* ≈ *N_t_T*/*C* in the linear approximation *N* ≈ *N_t_*, which holds when few of the FG motifs are bound. The total tethered Nup concentration, both free and bound, is *N_t_*. The fraction of transport factors that are bound is then given by

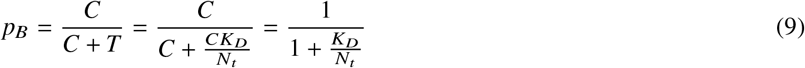

**Figure 2:**
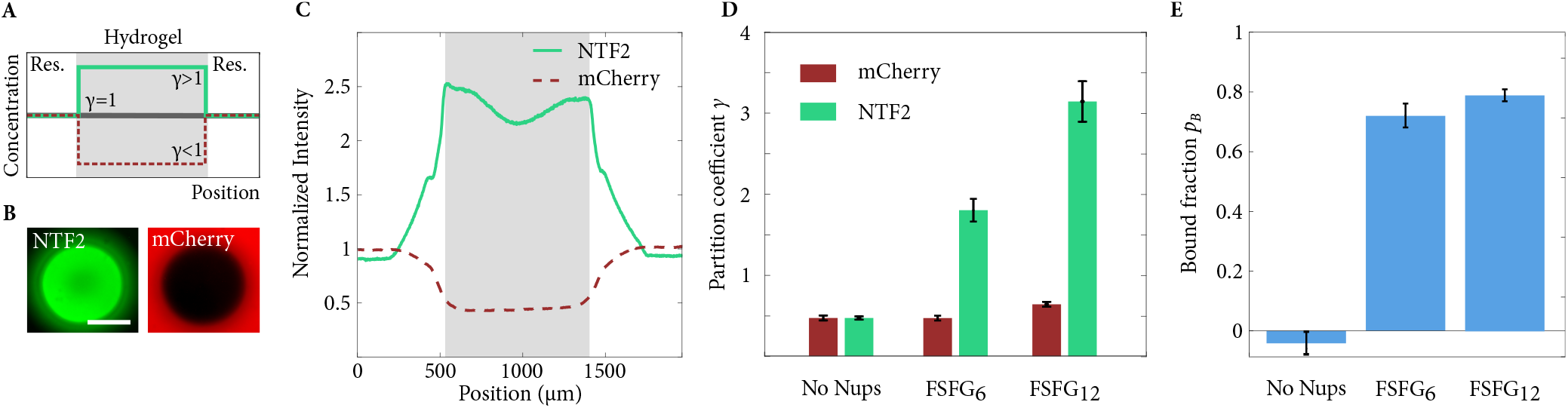
(A) Schematic of partition coefficient *γ* shown for a binding protein (*γ* > 1, green solid line) and non-binding protein (*γ* < 1, red dashed line). (B) Images of equilibrated gel and reservoir in NTF2 (left, green) and mCherry (right, red) channels. Scale bar is 500 *μ*m. (C) Fluorescence intensity profile of a gel containing FSFG_12_ and equilibrated with NTF2 and mCherry. Intensity profile is normalized to reservoir intensity. (D) Mean partition coefficients of NTF2 and mCherry for control, FSFG_6_, and FSFG_12_ gels. (E) Mean bound fraction *p_B_* for control, FSFG_6_, and FSFG_12_ gels. All error bars are standard error of the mean.

The concentrations of the inert protein and the transport factor in the reservoir are equal and given by *T*_0_. The total transport factor concentration within the gel is *T_t_* = *T* + *C* and is constant. If *γ*_*T*_ is the partition coefficient of the transport factor and *γ*_*I*_ that of the inert protein, then the transport factor concentration can be expressed as

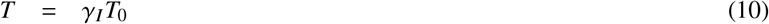

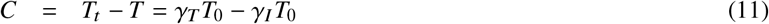

Therefore, within the gel, the chemical equilibrium condition can be expressed as

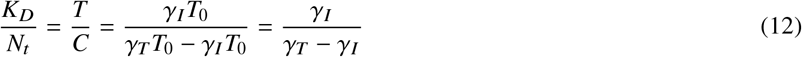

Combining Eqs. 9 and 12, the bound probability can be expressed in terms of the partition coefficients as

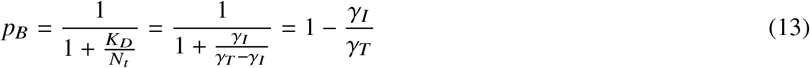

Therefore, we measured *p_B_* using the partition coefficients of the transport factor and inert protein, as determined from their equilibrated intensity within the gel as compared to that in the reservoir.

### Binding kinetics calculations

Comparison to theory requires an estimate of the off-rate *k*_off_ of the transport factor-Nup interaction. The off-rate calculation uses the relation between Nup concentration and dissociation constant described in the partition coefficient analysis:

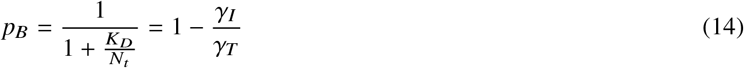

where *K_D_* = *k*_off_/*k*_on_, leading to

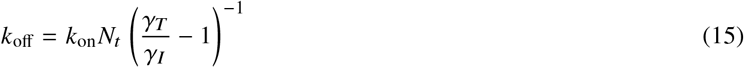

We assumed a diffusion-limited on-rate constant of *k*_on_ = 10^9^ s^−1^ M^−1^ (23, 38). We were unable to directly measure the tethered FG Nup concentration *N_t_*, but we were able to place upper and lower bounds on its value as described below. Using the upper bound of *N_t_* = 0.67 mM for both FSFG_6_ and FSFG_12_ gels, we calculated a dissociation constant of *K_D_* = 300 ± 80 *μ*M and an off-rate of *k*_off_ = 3.0 × 10^5^ ± 0.8 × 10^5^ *s*^−1^ for FSFG_6_. The corresponding values for FSFG_12_ are *K_D_* = 180 ± 20 *μ*M and *k*_off_ = 1.8 × 10^5^ ± 0.2 × 10^5^ *s*^−1^.

### Tether concentration and spacing

Knowledge of the tether concentration is necessary both for the binding kinetics calculation described above and to ensure that the contribution of inter-chain hopping to bound diffusion is minimized. Isolating the effect of tethered diffusion requires that transport factors be unlikely to interact with two tethers simultaneously, a condition which is met if the tether spacing is larger than the mean tether end-to-end distance. However, the sensitivity of our measurement also decreases with decreasing concentration. As a balance between these two effects, we designed our gels so that if all of the protein we introduced into the precursor solution is incorporated into the gels, the tether spacing would be at most equal to the mean end-to-end length as expected from a worm-like chain model.

To within the error of a BCA assay, the concentration of both FSFG_6_ and FSFG_12_ introduced into the gel is *N_t_* = 0.67 mM, giving an average spacing between tethers of 14 nm. To estimate the mean tether end-to-end length, we treated FSFG_6_ and FSFG_12_ as worm-like chains with persistence length *ℓ*_*p*_ = 1 nm and contour lengths of *L_c_* = 50 nm and 100 nm, respectively. The root-mean-squared end-to-end chain length is then given by (39)

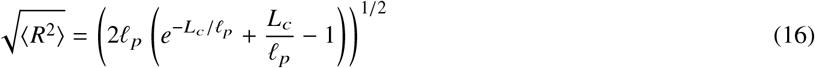

Using this model, FSFG_6_ is about 10 nm across and FSFG_12_ 14 nm.

We confirmed significant incorporation of the protein into the gel by 1) the binding of NTF2 to the gel, which predicts that all of the protein is incorporated and 2) by digesting the protein out of the gel using trypsin and measuring the resulting concentration in equilibrated solution. Note that the protein lacks any tyrosine and tryptophan residues, making only indirect techniques available. FG gels were soaked in a large volume of buffer to remove any free FSFG and then subjected to trypsin digestion (40, 41). The digested FSFG fragments were extracted and dried in a vacuum desiccator and their concentration quantified with a BCA assay. The final trypsin concentration of the sample was calculated to be approximately 10-fold below the BCA detection limit, a conclusion which was supported by the results of a trypsin-only sample in which no protein was detected. This assay resulted in a lower bound of 50 *μ*M (FSFG_6_) and 70 *μ*M (FSFG_12_) on the concentration of tethers, giving a mean spacing of 32 nm (FSFG_6_) and 29 nm (FSFG_12_). The mean spacing between tethers therefore falls between 14 nm (upper bound, *N_t_* = 0.67 mM) and 32 nm (lower bound, *N_t_* = 50 *μ*M) for FSFG_6_ and between 14 nm and 29 nm (*N_t_* = 70 *μ*M) for FSFG_12_, as compared to the 10 nm and 14 nm end-to-end distance of FSFG_6_ and FSFG_12_, respectively. Combined, these measurements indicate that tethered diffusion will be the predominant mechanism of bound diffusion, especially for FSFG_6_.

## RESULTS AND DISCUSSION

While previous work has shown that bound diffusion can lead to selective filtration (16–18), the molecular mechanisms that contribute to bound mobility and their relative importance remain poorly understood (13, 19). We focus on determining whether binding to flexible tethers can lead to appreciable bound motion. Transient binding to flexible molecular tethers is a feature of a number of biofilters (8, 42, 43), suggesting that tethered diffusion is a fundamental molecular mechanism that could be applicable in a variety of systems. Additionally, tethered diffusion could be used to design artificial selective filters.

In order to test the feasibility of tethered diffusion as a mechanism for bound mobility, we designed a material which displays tethered diffusion using a minimal set of biological components. We designed our material using key components of the NPC, FG Nups and a transport factor (Fig. 1), because the kinetics of the binding interaction are well established to be rapid, with diffusion limited on-rates (23, 38) and relatively weak avidity (24). We chose nuclear transport factor 2 (NTF2), a 28 kDa homodimer which binds to FG Nups (44, 45). As an inert comparison protein, we chose mCherry, a similarly-sized but non-binding red fluorescent protein. Our theory of tethered diffusion predicts that the bound diffusion constant should be affected by the length of the FG Nup fragment anchored to the gels. To quantitatively test the model’s predictions, we used as tethers two well-characterized peptides derived from the FG Nup Nsp1, FSFG_6_ (50 nm) and FSFG_12_ (100 nm) (Fig. 1A, Materials and Methods) (23, 24).

Several approaches can be used to measure bound diffusion. For example, both single-molecule and bulk measurements have been used to measure the bound diffusion of transcription factors (46–48). We chose to measure the bulk diffusion of particles within a uniform macroscopic material because the binding lifetime and distance traveled during a binding event are sufficiently short as to make single-molecule measurements difficult. Our bulk material consisted of polyacrylamide gels containing FG Nup fragments. A precursor solution containing acrylamide monomers, bisacrylamide crosslinkers, and bisacrylamide-labeled FSFG was added in microliter droplets to a flow chamber and polymerized through exposure to UV illumination (Fig. 1B). A solution containing fluorescent NTF2 and mCherry was introduced to the reservoir surrounding the gel (Fig. 1C). After a 24-hour equilibration period, FRAP experiments were performed on the gels. In control gels in the absence of FG Nups, NTF2 and mCherry showed similar behavior, and gels made from multiple protein purifications and conjugation reactions gave consistent results.

Our experiments were designed to isolate tethered diffusion and to avoid other possible sources of bound diffusion. These could include other features known to be important to nuclear transport, such as FG Nup cohesiveness (49, 50), active release of cargo and transport factors from the pore (10, 51, 52), and contributions of crowding and transport factors themselves to maintaining the selective barrier (53, 54). To retain only the features of the NPC which contribute to tethered diffusion, we chose an FG Nup fragment which is not cohesive (55) and for which we saw no aggregation or liquid phase separation at concentrations up to 20 mg/mL. We omitted proteins which could contribute to active release (*e.g.*, Ran). Multivalent FG Nup-transport factor interactions could permit transport factors to move between nearby Nups without fully unbinding (14, 15) or to crosslink adjacent FG Nups, altering the barrier properties of the gel (53). These effects should only be important to bound-state diffusion if the density of FG Nups within the gel is large enough that a transport factor is likely interact with two FG Nups at once. By tuning the average concentration of tethered FG Nups within the gels, we developed gels in which the mean tethered peptide spacing is larger than its end-to-end distance (see Materials and Methods). Therefore, the predominant mechanism of bound-state mobility in these gels should be tethered diffusion.

To determine the bound diffusion constant of a protein within an FG gel, we require knowledge only of the fraction of time the protein spends bound to FSFG and its diffusion constant arising from both bound and free motion. The bound fraction can be calculated from the partition coefficient. A protein’s partition coefficient *γ* is the ratio of its concentration within an equilibrated gel to that in the surrounding reservoir (Fig. 2A). The partition coefficients were measured in equilibrated gels by comparing the intensity within the gel to that of the reservoir. NTF2 and mCherry partitioned similarly into gels that contained no FG Nups, indicating that both proteins interact similarly with the inert polyacrylamide gel scaffold (Fig. 2D). The mCherry partition coefficient remained nearly the same in both FG-containing gels as in the control gels without FG Nups. This indicates that the FG Nup neither occupies a signifiant fraction of the gel volume nor significantly alters the gel properties. The partition coefficient of NTF2 increased dramatically in the presence of FSFG_6_ and FSFG_12_, consistent with binding. The fraction bound is given by the the selective enrichment of NTF2 relative to mCherry in the gels due to binding (Eq. 13, Fig. 2E).

We used FRAP to measure the diffusion constant of mCherry and NTF2 within the gels. FRAP relies on the gradual redistribution of fluorescent transport factors after a region of the gel is photobleached; the recovery time of the bleached region is related to the transport factor’s diffusion constant (31). In some cases, the recovery time can be well fit by simple models of diffusion and binding. However, in our experiments there was significant exchange between the gel and the reservoir, and the gels were not always fully equilibrated. To accurately measure the diffusion constant with these confounding effects, we used a time-dependent, two-dimensional Fourier series solution to the diffusion equation (37). The two-dimensional post-bleach concentration profile was taken as the initial condition. We calculated the appropriate Fourier coefficients and then simulated the time-dependent recovery of the bleached region. This approach gave reconstructed images and recovery curves consistent with those observed experimentally (Fig. 3A, B). The diffusion constant was extracted from the fits for both NTF2 and mCherry in all three gel conditions (Fig. 3C). The diffusion constants for the transport factor and inert protein were roughly equal in the control gels, indicating that there is no significant difference between the proteins in their interaction with the polyacrylamide scaffold (*p* = 0.61). The transport factor, NTF2, had a lower diffusion constant in the gels containing FG Nups, as expected, since motion is slowed during binding events.

**Figure 3:**
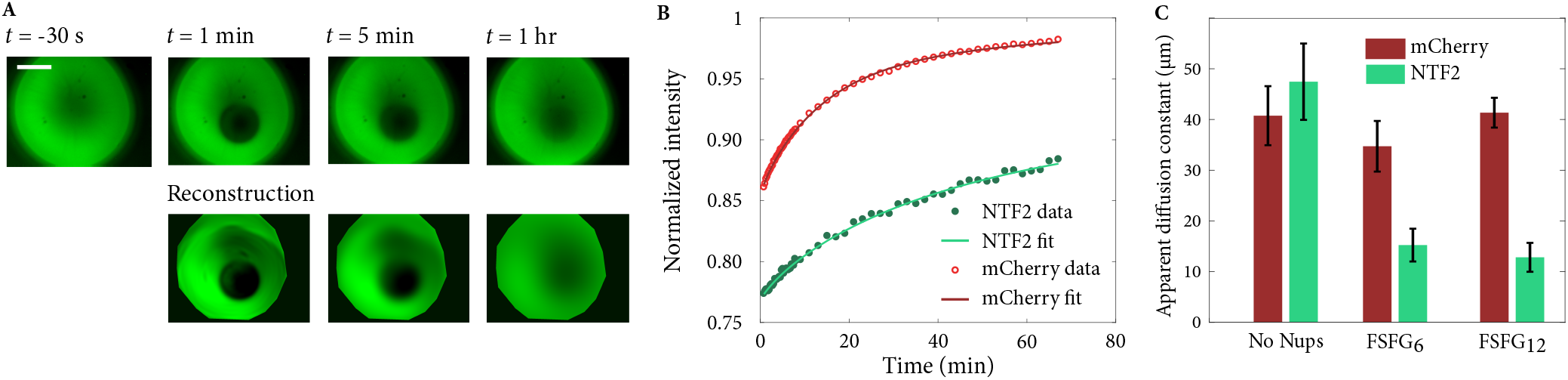
(A) Bleaching and recovery of a gel containing FSFG_6_. NTF2 channel shown only. Comparison to simulated data shown. Scale bar is 500 *μ*m. (B) Normalized recovery curves and Fourier series fit for gel shown in (A). (C) Diffusion constant for NTF2 and mCherry in each experimental condition. Error bars are standard error of the mean.

To determine the bound diffusion constant, we assumed that the diffusion constant *D* of NTF2 is given by a weighted average of the free and bound diffusion constants since NTF2-FSFG binding is fast relative to the diffusion of NTF2. The diffusion coefficient is given by

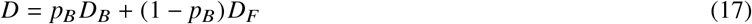

where *D_B_* is the bound diffusion constant of NTF2, *D_F_* is its free diffusion constant (while unbound), and *p_B_* is the fraction of time that NTF2 spends bound. We take the free diffusion constant of the transport factor to be equal to the diffusion of the inert protein mCherry.

We measured significant bound diffusion for NTF2 in both FG-Nup-containing gels (Fig. 4). The bound diffusion constant of NTF2 in the FSFG_6_ gels was 5.6 ± 2.2 *μ*m^2^/s (*D_B_*/*D_F_* = 0.24 ± 0.09), and in the FSFG_12_ gels it was 5.2 ± 3.2 *μ*m^2^/s (*D_B_*/*D_F_* = 0.13 ± 0.08). These results demonstrate that biomaterials which feature transient binding to dynamic polymers can display bound-state diffusion.

**Figure 4:**
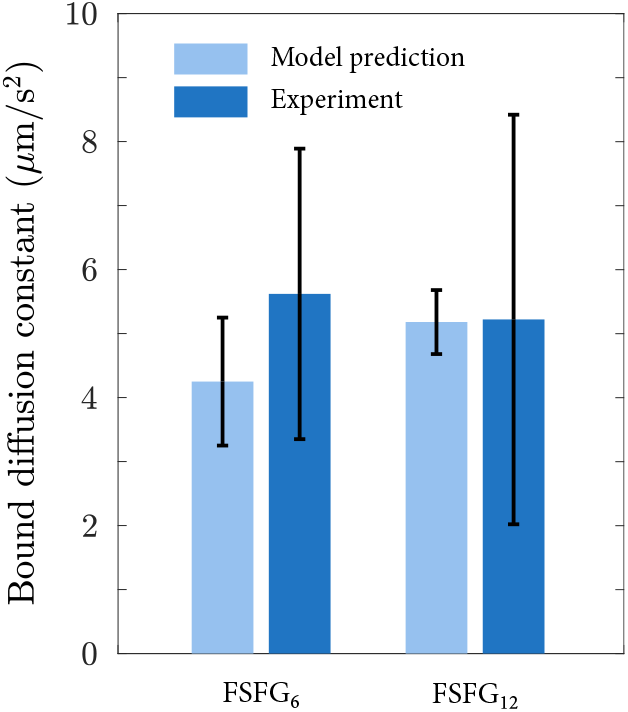
Model predictions and experimental measurements of the bound diffusion constant *D_B_* for NTF2 in the FSFG_6_ and FSFG_12_ gels. The tethered diffusion model (Eq. 18) was used for the predictions, assuming a tether concentration of 0.67 mM in both conditions and a diffusion-limited on-rate constant of 10^9^ M^−1^ s^−1^. The predicted value of *D_B_* was computed for each gel and averaged for each condition. Error bars are standard error of the mean.

We previously predicted the expected bound diffusion constant arising from tethered diffusion (13). In this theory, the protein was modeled as diffusing in a harmonic potential well representing the tether to which it is bound. The bound diffusion is determined by the mean-squared displacement of a particle during a binding event weighted by the probability of that binding lifetime. The resulting bound diffusion constant is given by

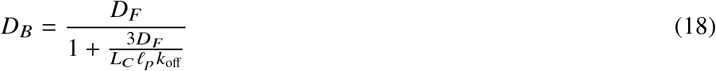

where *L_C_* is the tether contour length, *ℓ*_*p*_ is its persistence length, and *k*_off_ is the off-rate of the interaction between the transport factor and FG Nup. The persistence length of disordered proteins is about 1 nm (56), and the tether contour length is ~50 nm for FSFG_6_ and ~100 nm for FSFG_12_. The off-rate, *k*_off_, was estimated using the maximum possible tethered FG Nup concentration (see Materials and Methods) and a diffusion limited on-rate (23, 38). We calculate a mean maximum dissociation constant of *K_D_* = 24 ± 6 *μ*M for FSFG_6_ and *K_D_* = 304 ± 78 *μ*M for FSFG_12_. These values are significantly lower (tighter) than those recently measured by NMR and isothermal titration calorimetry (24), which may be due to the crowded gel environment. With these assumptions, the predicted value of *D_B_* was calculated for each gel using Eq. 18 and then averaged (Fig. 4).

For both FSFG_6_ and FSFG_12_, the measured value of bound diffusion constant is consistent with the value predicted by our minimal model of tethered diffusion. Although FSFG_12_ is twice as long as FSFG_6_, the affinity is also weaker. As the bound diffusion constant only depends on the product *L*_*C*_*ℓ*_*p*_*k*_off_, the bound diffusion constant in the FSFG_12_ gel is only expected to be slightly higher than that in the FSFG_6_ gel (Fig. 4).

## CONCLUSION

Biological filters are necessary for cells and organisms to control the localization of macromolecules. They are typically made of flexible polymers which provide a selective barrier, allowing the passage of some macromolecules while inhibiting the passage of others (8, 42, 43). Bound-state diffusion is necessary for the selective enhancement of the steady-state flux for binding particles relative to similar inert particles (13, 16–18). In the case of the NPC, those particles which are able to passage the barrier often have relatively weak affinities and rapid binding dynamics (23, 38). The combination of binding to sites on flexible polymers with rapid binding dynamics is thought to be sufficient for significant diffusion even within the bound state, an effect termed tethered diffusion (13).

In order to experimentally determine whether bio-inspired gels could be designed to utilize tethered diffusion, we tethered FG Nup fragments to polyacrylamide gels and measured the diffusion and binding properties of a transport factor (NTF2) and comparable inert protein (mCherry) (Fig. 1). We designed our gels to allow for accurate measurement of the partition coefficient (and thus the fraction of time bound, Fig. 2) and diffusion constant (Fig. 3), the two parameters needed to determine the bound diffusion. By design, the polyacrylamide gel scaffold interacts similarly with both test proteins, leading to diffusion constants and partition coefficients that are similar in control gels lacking FG Nups. Binding increases the concentration of NTF2 in the gel and slows its motion. Bound diffusion contributes significantly to the motion of NTF2 in FG-containing gels; NTF2 motion would be reduced by over 10% in its absence. The bound diffusion constants of NTF2 calculated using the diffusion constants obtained by FRAP are quantitatively consistent with theoretical predictions of diffusion while bound to flexible tethers (Fig. 4) (13).

These results represent a first step toward understanding and utilizing tethered diffusion in biological and bio-inspired systems. While our system utilizes components of cells’ nuclear transport machinery, the requirements of rapid binding kinetics and binding site flexibility are present in other systems. Tethered diffusion may be used to engineer artificial bio-inspired filters or to manipulate the flux of particles through naturally-occurring filters such as mucus membranes. Binding interactions can be used to impart selectivity, while tethered diffusion would allow for particle motion necessary for the particle to penetrate into and across a barrier.

## AUTHOR CONTRIBUTIONS

L.E.H., M.D.B., and L.M. designed the research and wrote the article. L.M. carried out the experiments and analyzed the data.

## ACKNOWLEDGMENTS

The authors would like to thank Joseph Dragavon (BioFrontiers Advanced Light Microscopy Core) for assistance with microscopy and Eric Verbeke, Sadhana Sharma, Stephanie Bryant, Benjamin Fairbanks, and Nathan Crossette for assistance with gel design and fabrication. This work is supported by NIH R35 GM119755, NSF DMR-1551095, NSF MRSEC DMR-1420736, the Boettcher Foundation, and a CU innovative seed grant.

